# Circadian Control of Sleep by Melatonin via MT_1_-Dependent Activation of BK Channels in the Suprachiasmatic Nucleus

**DOI:** 10.1101/2025.03.12.642893

**Authors:** Kiranmayi Vedantham, Adeel Ahmad, Longgang Niu, Yuan Shui, Fouad Lemtiri-Chlieh, Deborah Kaback, Xin-Ming Ma, Xiangyou Hu, Siu-Pok Yee, Zhao-Wen Wang

## Abstract

Melatonin promotes sleep through mechanisms that have remained elusive. Here, we identify a molecular pathway by which melatonin promotes sleep by activating BK channels (Slo1) via MT_1_ receptors in the suprachiasmatic nucleus (SCN), the brain’s master circadian clock. In melatonin-proficient CBA/CaJ mice, knockout of either *MT_1_* or *Slo1* reduces REM and NREM sleep during the rest phase (daytime), accompanied by prolonged action potentials and diminished afterhyperpolarization in SCN neurons. These electrophysiological and behavioral changes are minimal during the active phase (nighttime). Strikingly, Slo1 expression in the SCN peaks during the daytime, contrary to previous reports, but aligning with its sleep-promoting function. *Slo1*, but not *MT_1_*, deletion also triggers spontaneous seizures, highlighting broader functions beyond circadian control. Structural mapping identifies critical domains mediating MT_1_-Slo1 coupling. Together, these findings position the MT_1_-Slo1 signaling axis as a core circadian mechanism linking melatonin to sleep regulation and a potential therapeutic target for sleep disorders.

## INTRODUCTION

Melatonin, often referred to as the “dark hormone”, is a key regulator of circadian rhythms and sleep. It exerts its effects primarily through two G protein-coupled receptors, MT_1_ and MT_2_ ^1^. However, the downstream effectors that mediate melatonin’s sleep-promoting actions in mammals have not been identified.

Knockout mouse studies have implicated MT_1_ melatonin receptors and BK channels (Slo1)— large-conductance, Ca^2+^- and voltage-activated K^+^ channels—in circadian rhythm regulation. In the melatonin-proficient C3H/He strain, MT_1_ deletion significantly reduces REM sleep, primarily during the subjective daytime (inactive phase) ^2^. In the melatonin-deficient C57BL/6 strain, MT_1_ deletion prevents the circadian phase advance of wheel-running activity that melatonin treatment induces in wild-type mice ^3^. A melatonin analog also more effectively phase-advances electrical activity in slices of the suprachiasmatic nucleus (SCN) from wild-type than *MT_1_* knockout mice of the same strain ^4^. Separately, *Slo1* knockout impairs circadian locomotor rhythmicity in wheel-running assays in a hybrid mouse strain that lacks melatonin secretion ^5^. Despite these findings, the specific contributions of MT_1_ and Slo1 to sleep regulation remain unclear.

The SCN of the hypothalamus functions as the central circadian clock, synchronizing behavioral and physiological rhythms, including sleep-wake cycle, with the environmental light-dark cycle via direct retinal input ^6–10^. Both MT_1_ receptors ^11–14^ and Slo1 are expressed in the SCN. In SCN brain slices from wild-type, melatonin application reduces neuronal firing, an effect absent in slices from *MT_1_* knockout mice ^4^; however, this result was not replicated in a separate study ^3^. Genetic ablation of *Slo1* increases SCN firing rates at night ^5^, and deletion of the Slo1 β2-subunit disrupts daily firing rhythmicity ^15^. These findings suggest that both MT_1_ and Slo1 contribute to the modulation of SCN neuronal excitability. Nonetheless, whether MT_1_ and Slo1 functionally interact within the SCN and whether such interactions regulate sleep remain unknown.

In *C. elegans*, a period of quiescence between consecutive developmental stages, commonly referred to as lethargus or developmentally timed quiescence (DTQ), exhibits behavioral similarities to sleep in mammals ^16^. We recently discovered that melatonin increases the duration of DTQ by activating SLO-1 (a mammalian Slo1 ortholog) through a specific melatonin receptor (PCDR-1), and that genetic deletion of either *slo-1*, *pcdr-1*, or an enzyme required for melatonin synthesis (*homt-1*) leads to shortened DTQ duration ^17^. In addition, we found that melatonin can activate human Slo1 in *Xenopus* oocytes in the presence of MT_1_ but not MT_2_ receptors ^17^. These findings suggest that Slo1 might represent a conserved molecular target through which melatonin regulates quiescence, potentially relevant to sleep in mammals. However, this hypothesis requires direct testing in mammals, as *C. elegans* DTQ diverges from mammalian sleep in critical ways: it is developmentally rather than circadian-timed, and lacks conserved neural circuitry (e.g., hypothalamic nuclei) and most mammalian clock genes.

In this study, we examined the roles of MT_1_ and Slo1 in regulating SCN neuronal electrical properties and sleep behavior in CBA/CaJ mice, a melatonin-proficient strain ^18–20^. Using whole-cell patch-clamp recordings, EEG/EMG monitoring, and coimmunoprecipitation assays, we identified an MT_1_-Slo1 signaling axis that promotes sleep by modulating action potential (AP) properties in SCN neurons. Importantly, we show that MT_1_ and Slo1 regulate AP waveform features during the subjective day without affecting firing frequency, challenging previous studies that emphasized Slo1’s role in nocturnal firing rhythms ^5^. Additionally, we observed pronounced seizure activity in *Slo1* knockout mice, consistent with growing evidence linking Slo1 dysfunction to epilepsy in humans ^21,22^.

Together, this study reveals that MT_1_ and Slo1 constitute a molecular pathway by which melatonin promotes sleep in mammals. Our findings redefine the circadian functions of MT_1_ and Slo1, positioning them as critical regulators of daytime sleep maintenance and as promising therapeutic targets for sleep disorders. Moreover, we establish Slo1 knockout mice as a valuable model for studying epilepsy related to Slo1 dysfunction.

## RESULTS

### Generation of *MT_1_* and *Slo1* knockout mice

We used the CRISPR-Cas9 strategy to generate knockout strains for *Mtnr1a* and *Kcnma1*, referred to as *MT_1_^-/-^*and *Slo1^-/-^*, respectively. Littermate controls are designated as *MT_1_^+/+^* and *Slo1^+/+^*.

The *MT_1_^-/-^* strain carries a 500-bp deletion that eliminates the entire first exon, along with adjacent promoter and intronic regions (Fig. 1a-c and Supplementary Fig. 1a, b). The removal of exon 1’s coding sequence and the splice donor site for exon 2 (the gene has only two exons) renders this mutation null, preventing the production of any MT_1_ protein.

**Fig. 1:**
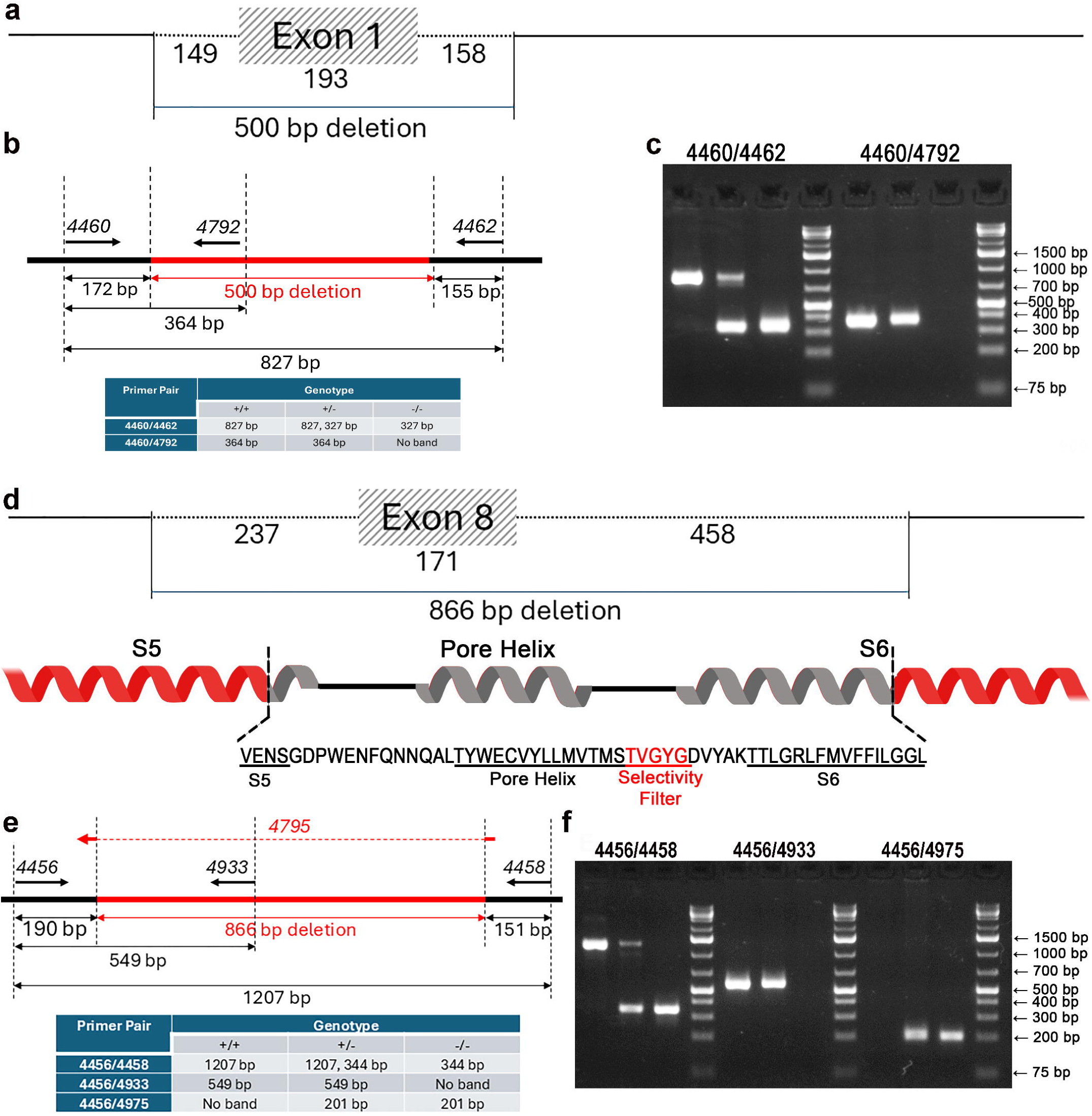
Generation of *MT_1_^-/-^*and *Slo1^-/-^* mice. **a** Schematic representation of the *Mtnr1a* (*MT_1_*) gene, indicating the location and size of the deletion. **b** Schematic diagram of PCR primers used for genotyping, with the expected PCR product sizes for *MT_1_^+/+^, MT_1_^+/-^ and MT_1_^-/-^* provided in the table. **c** Gel image showing PCR genotyping results for *MT_1_^+/+^, MT_1_^+/-^ and MT_1_^-/-^* mice. **d** Schematic representation of the *Kcnma1* (*Slo1*) gene, indicating the location and size of the deletion, and an α-helical representation of Slo1 highlighting the deleted amino acids within the core structure of the channel. **e** Schematic diagram of PCR primers used for genotyping, with the expected PCR product sizes for *Slo1^+/+^, Slo1^+/-^ and Slo1^-/-^* provided in the table. **f** Gel image showing PCR genotyping results for *Slo1^+/+^, Slo1^+/-^ and Slo1^-/-^* mice.

The *Slo1^-/-^* strain has an 866-bp deletion (and 3 bp insertion) that removes the entire exon 8 (as designated in *Kcnma1* isoform 203), along with portions of the adjacent introns 7 and 8 (Fig. 1d-f and Supplementary Fig. 1c, d). Exon 8 is conserved across all *Kcnma1* isoforms and encodes a critical region of the Slo1 channel, including 4 residues in the S5 transmembrane segment, the channel pore, and 16 residues in the S6 transmembrane segment (Fig. 1d). This in-frame deletion abolishes K^+^ conductance but may still allow the production of a defective Slo1 protein.

### *Knockout of Slo1* or *MT_1_* reduces sleep duration

We investigated the effects of *MT_1_^-/-^* and *Slo1^-/-^* on sleep-wake behavior by performing EEG and EMG recordings on mice under regular light/dark cycle (12:12). Periods of wakefulness, NREM sleep, and REM sleep were identified by analyzing EEG and EMG traces. Wakefulness period was characterized by low-amplitude EEG and high-amplitude EMG; NREM period by high-amplitude EEG with a dominant frequency of < 4 Hz and low-amplitude EMG; and REM period by low- or high-amplitude EEG with a dominant frequency between 4 and 8 Hz and low-amplitude EMG (Fig. 2a).

**Fig. 2.**
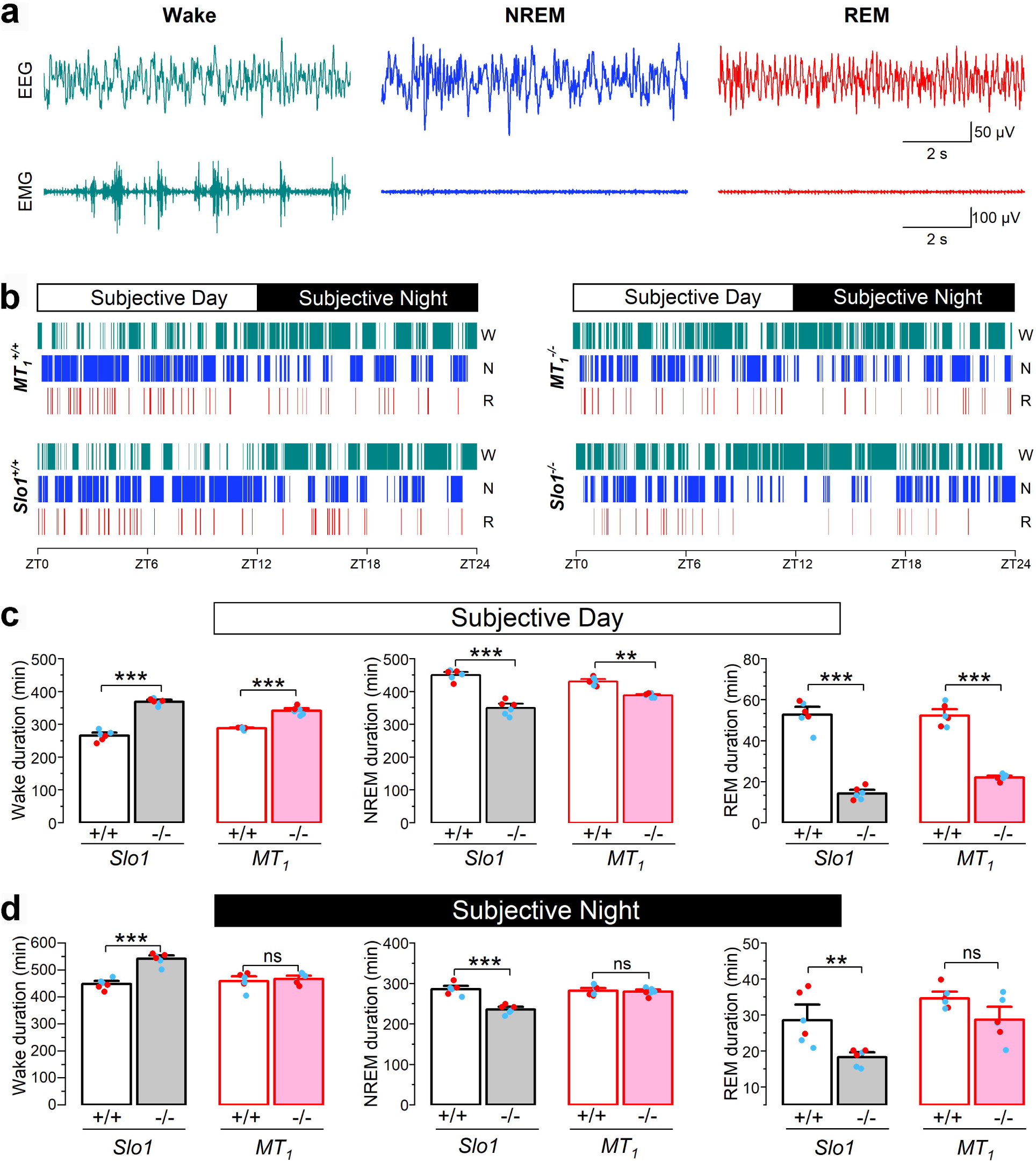
*Slo1^-/-^*and *MT_1_^-/-^* mice exhibit reduced REM and NREM sleep and increased wakefulness primarily during the subjective daytime. **a** Representative traces of electroencephalogram (EEG) and electromyogram (EMG) showing distinguishing features of REM sleep, NREM sleep, and wakefulness. **b** Representative hypnograms derived from EEG and EMG recordings for *Slo1^+/+^*, *Slo1^-/-^*, *MT_1_^+/+^*, and *MT_1_^-/-^* mice. **c** Quantification of sleep and wake durations during the subjective daytime. **d** Quantification of sleep and wake durations during the subjective nighttime. In panels **c** and **d**, the sample size (*n*) was 6 per group, including three males (blue data points) and three females (red data points). Asterisks indicate statistically significant differences between knockout mice and their corresponding littermate controls (* *p* < 0.05; ** *p* < 0.01; *** *p* < 0.001), while pound symbols indicate significant differences between *Slo1^+/+^* and *MT_1_^+/+^*or between *Slo1^-/-^* and *MT_1_^-/-^* (# *p* < 0.05; ## *p* < 0.01). Statistical analyses were performed using one-way *ANOVA* followed by Tukey’s post hoc test.

During the subjective day (light on), both *MT_1_^-/-^*and *Slo1^-/-^* mice exhibited significantly increased wake time and decreased NREM and REM durations, with an apparently greater reduction in REM than NREM (Fig. 2b, c). The magnitudes of changes in wake, NREM, and REM durations caused by *MT_1_^-/-^* and *Slo1^-/-^* mutations, compared to their respective littermate controls, averaged 18.8% and 39.2%, 9.5% and 20.1%, and 59.6% and 73.5%, respectively. During the subjective night (light off), the *MT_1_^-/-^* mutation had no significant effect on wake or sleep durations, whereas *Slo1^-/-^*mice showed significant decreases in both NREM and REM sleep durations and a significant increase in wake time compared to littermate controls (Fig. 2b, d). The distribution patterns of male and female data points in the bar graphs (Fig. 2c, d) suggest that *MT_1_^-/-^*and *Slo1^-/-^* exert similar effects on sleep behavior in male and female mice.

### *Knockout of Slo1, but not MT_1_*, causes significant ictal EEG activity

Our analysis of EEG data revealed frequent high-amplitude, rhythmic electrical discharges in *Slo1^-/-^* mice, characteristic of seizure activity (Fig. 3a). These discharges occurred exclusively between Zeitgeber time (ZT) 5 and ZT 21 and were not observed in any of the other mouse strains (*Slo1^+/+^*, *MT_1_^+/+^*, and *MT_1_^-/-^*).

**Fig. 3:**
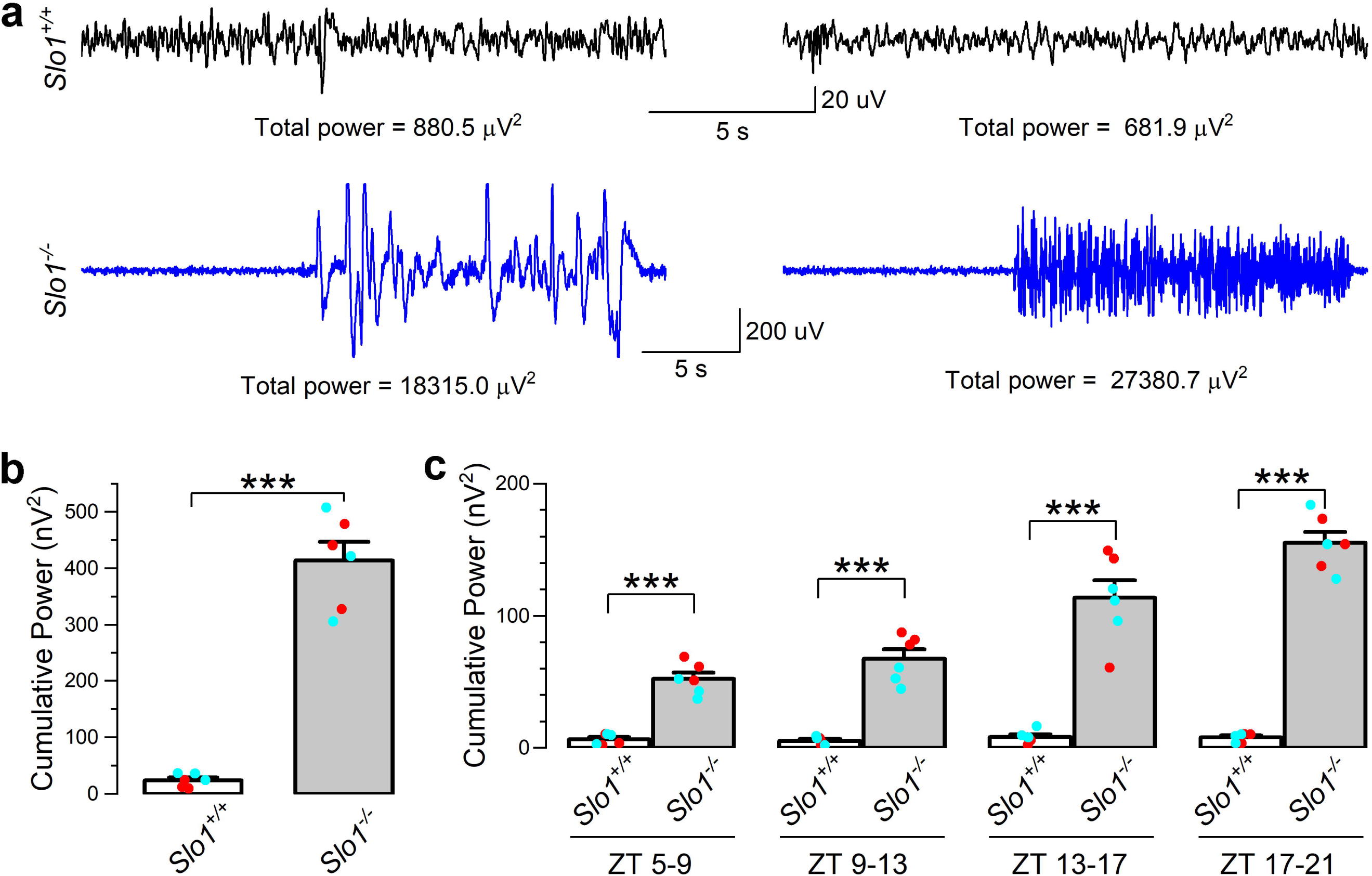
*Slo1^-/-^*mice exhibit seizure activity. **a** Representative electroencephalogram (EEG) traces from *Slo1^+/+^* and *Slo1^-/-^* mice. **b** Comparison of cumulative EEG power between *Slo1^+/+^* and *Slo1*^⁻^*^/^*^⁻^ mice. For *Slo1^-/-^* mice, cumulative EEG power was calculated as the sum of the total power of all seizure events. For *Slo1^+/+^* mice, which exhibited no seizure activity, cumulative EEG power was calculated as the sum of the total power of randomly selected EEG segments. These segments matched the mean number (18) and mean duration (17.6 seconds) of seizure events observed in the *Slo1*^⁻^*^/^*^⁻^ group. **c** Comparison of cumulative EEG power between Slo1+/+ and Slo1⁻/⁻ mice, analyzed based on Zeitgeber time (ZT) and grouped into 4-hour intervals. In **b** and **c**, the sample size (*n*) was 6 per genotype, including three males (blue data points) and three females (red data points). Statistical significance (***, p < 0.001) between Slo1⁻/⁻ mice and their littermate controls was determined using an unpaired t-test in **b** and a two-way mixed-design ANOVA followed by Tukey’s post hoc test in **c**.

We quantified the number and mean duration of ictal events, the total power of each event, and the cumulative power of all ictal events in each mouse. The mean ictal event number between ZT 5 and ZT 21 in *Slo1^-/-^* mice was 17.5 ± 1.8, with a mean duration of 17.6 ± 1.4 seconds.

To compare power levels, we randomly selected 18 EEG segments from each *Slo1^+/+^* mouse, with each being 17.6 seconds long to match the number and duration of ictal events in *Slo1^-/-^* mice. The cumulative power in these segments from wild-type mice was then compared to that of *Slo1^-/-^* mice, revealing a more than 40-fold increase in cumulative power in *Slo1^-/-^*mice (Fig. 3b).

Additionally, we divided the cumulative power into four equal time periods (4 hours each) and compared them to those of *Slo1^+/+^*mice. This analysis showed that ictal activity in *Slo1^-/-^* mice increased gradually over the successive periods (Fig. 3c).

Together, our analysis indicates that *Slo1^-/-^* leads to strong seizure activity occurring within a specific time window of the day. Since we quantified seizure activity only for ictal events lasting ≥10 seconds, the actual magnitude of the difference between *Slo1^+/+^* and *Slo1^-/-^* mice is larger than shown in the figure.

Notably, the periods of high seizure activity (occurring between ZT 5 and ZT 21 largely overlapped with the time window when *Slo1^-/-^*mice, but not *MT_1_^-/-^* mice, exhibited decreased REM and NREM sleep and increased wakefulness. This temporal correlation suggests that the increased seizure activity in *Slo1^-/-^* mice contributed to the observed alterations in sleep and wake behaviors.

### *Knockout of Slo1* or *MT_1_* broadens APs and suppresses AHP during the daytime

To determine where MT_1_ and Slo1 might function to regulate sleep, we analyzed the effects of *Slo1^-/-^* and *MT_1_^-/-^*on SCN neuronal electrical properties. Given that both *Slo1^-/-^*and *MT_1_^-/-^* exhibited differing effects on sleep between the daytime and nighttime, we investigated their effects on SCN neurons in both the subjective day and night. Daytime effects were assessed using mice housed under a light cycle with lights on from 7:00 am to 7:00 pm and off from 7:00 pm to 7:00 am. Brain slices were prepared between 10:00 and 11:00 am and used for electrophysiological experiments between 2:00 and 5:00 pm, corresponding to ZT 3-4 and ZT 7–10, respectively.

We performed whole-cell patch-clamp experiments on neurons located in the ventromedial SCN. Our daytime experiments utilized *Slo1^-/-^* and *MT_1_^-/-^* single knockout strains, their respective littermate controls, and a *Slo1^-/-^;MT_1_^-/-^*double knockout strain.

There were no significant differences in membrane resistance (R_m_) or resting membrane potential (RPM) between the different groups (Fig. 4a). AP frequencies in response to current injections (-10 pA to 20 pA in 5-pA increments, with a pulse duration of 5 seconds) tended to increase with higher current injections; however, there was no significant difference between the different groups at each current injection step (Fig. 4b), suggesting that neither Slo1 nor MT_1_ significantly influences AP frequency.

**Fig. 4.**
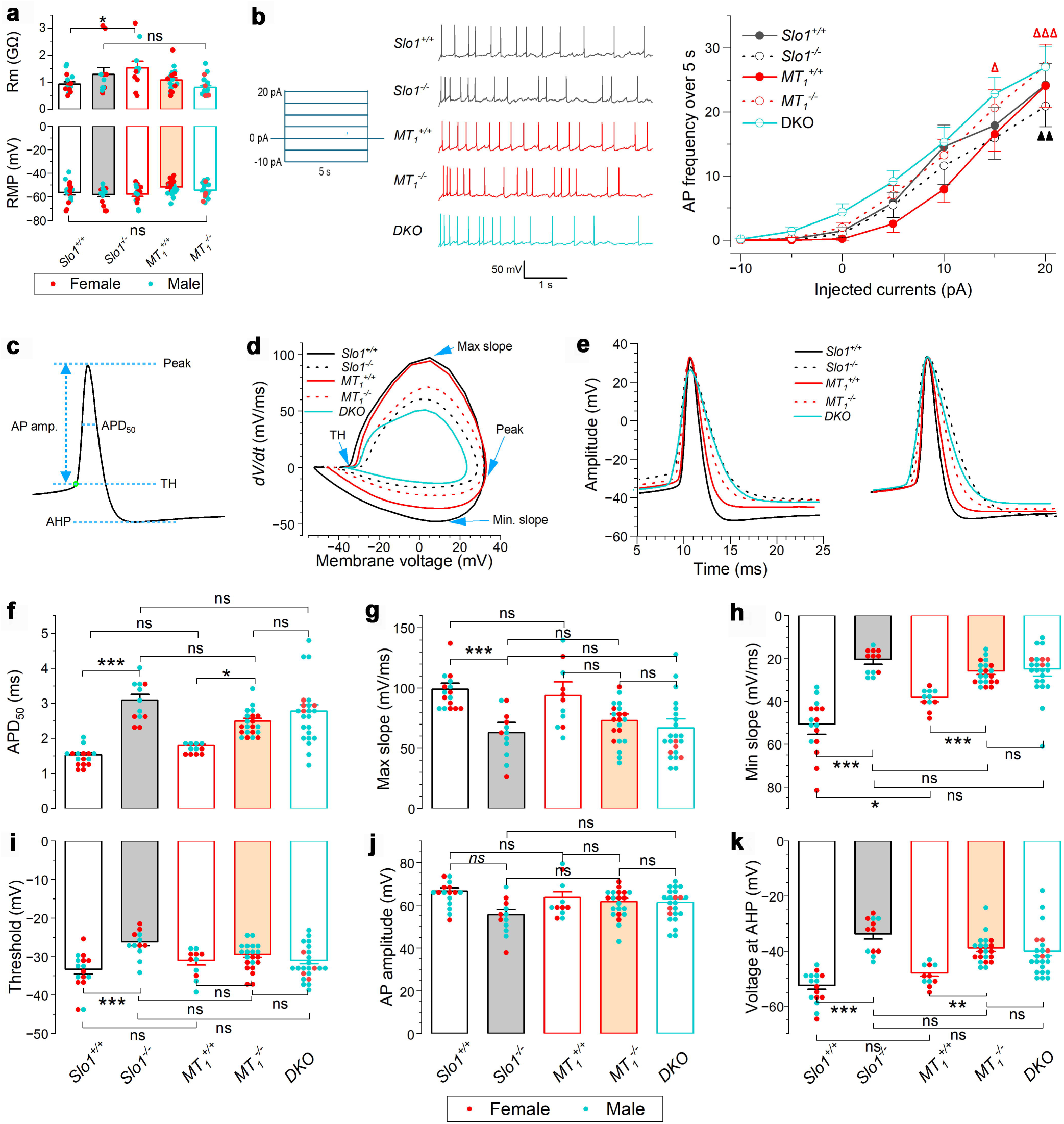
SCN neurons in *Slo1^-/-^*and *MT_1_^-/-^* mice exhibit prolonged action potential (AP) duration and suppressed afterhyperpolarization (AHP) during the subjective day. **a** Membrane resistance (*R_m_*) and resting membrane potential (RMP) are unaffected by *Slo1^-/-^* or *MT_1_^-/-^.* b *Slo1^-/-^* or *MT_1_^-/-^* does not alter the frequency of APs induced by current injections. *Left*: Diagram of the current injection protocol. *Middle*: Representative current-clamp traces during 15-pA current injection. *Right*: AP frequency as a function of current injection. Filled and open triangles indicate statistically significant differences compared to the AP frequency at 0-pA current injection for the same group (Δ *p* < 0.05; ΔΔ *p* < 0.01; ΔΔΔ *p* < 0.001). Statistical analyses were performed using two-way mixed-design *ANOVA* followed by Tukey’s post hoc test. **c** Representative AP trace illustrating the quantification of AP threshold (TH), AP amplitude, APD_50_ (AP duration at 50% repolarization), and AHP. **d** Voltage phase plots of group averaged APs induced by 15-pA current injection. Blue arrows indicate TH, AP peak, and maximum and minimum slopes. **e** Group-averaged APs induced by 15-pA current injection, displayed as actual (left) and normalized (right) waveforms. **f-k** Quantitative comparisons of AP-related parameters. Asterisks indicate statistically significant differences between groups (* *p* < 0.05; ** *p* < 0.01; *** *p* < 0.001), while “ns” indicates no significant difference. Statistical analyses were performed using two-way *ANOVA* (factors: genotype and time of day) followed by Tukey’s post hoc test. The number of neurons recorded (*n*) were: *Slo1^+/+^* 16, *Slo1^-/-^*12, *MT_1_^+/+^* 11, *MT_1_^-/-^*21, and *MT_1_^-/-^ ;Slo1^-/-^* double knockout (DKO) 23.

We selected APs induced by 15- and 20-pA current injections to quantify several AP-related parameters, including AP duration at 50% repolarization (APD_50_), maximum and minimum slopes, threshold, amplitude, and afterhyperpolarization (AHP), with the results shown in Fig. 4c-k and Supplementary Fig. 2a-h, respectively. Both *Slo1^-/-^*and *MT_1_^-/-^* mice showed a substantial increase in APD_50_, and this phenotype was not aggravated in *Slo1* and *MT_1_*double knockout mice (Fig. 4e, f; Supplementary Fig. 2b, c). The prolongation of APD_50_ in the knockout mice compared to littermate controls was mainly due to a much slower rate of repolarization, as indicated by the group-averaged AP waveforms (Fig. 4e; Supplementary Fig. 2b) and the AP minimum slopes (Fig. 4h; Supplementary Fig. 2d, e). *Slo1^-/-^*mice also exhibited significant decreases in the AP maximum slope and threshold, which were not observed in *MT_1_^-/-^* mice (Fig. 4g, i and Supplementary Fig. 2d, f). These results indicate that both Slo1 and MT_1_ are crucial for AP repolarization, and a loss of their function significantly prolongs AP duration. The differential effects between *Slo1^-/-^* and *MT1^-/-^*on AP maximum slope and threshold suggest that Slo1 function is not entirely dependent on MT_1_.

Knockout mice also exhibited a markedly suppressed AHP, defined as the hyperpolarized membrane voltage level relative to the threshold following each AP in *Slo1^+/+^* and *MT_1_^+/+^*mice. While AHP was obvious in littermate controls and could be automatically detected (reported as the membrane voltage level) using our analysis software, it was absent in *Slo1^-/-^*and *MT_1_^-/-^* mice (Fig. 4e; Supplementary Fig. 2b). To allow quantitative comparisons between the knockout mice and their littermate controls, we manually measured the membrane potential at a post-AP peak time point (5 ms) in *Slo1^-/-^* and *MT_1_^-/-^* mice, corresponding to the mean interval between the AP peak and automatically detected AHP in *Slo1^+/+^* mice (4.7 ms). The membrane potential in knockout mice was substantially more depolarized than the AHP in their littermate controls (Fig. 4k, Supplementary Fig. 2h). This analysis underscores the essential roles of both Slo1 and MT_1_ in AHP generation.

Male and female knockout mice exhibited qualitatively similar changes in the quantified AP parameters compared to littermate controls. They differed significantly in the mean values of a few measured parameters, although these differences were inconsistent between the 15-pA and 20-pA current injection steps (Supplementary Fig. 3). Overall, these findings suggest that there are no major sex-specific differences in the effects of *MT_1_* and *Slo1* knockout on AP-related parameters.

### *Knockout of Slo1* or *MT_1_ does not affect* AP duration and AHP during the nighttime

To evaluate the nighttime effects of *Slo1 or MT_1_* knockout on SCN neurons, we performed electrophysiological experiments using mice housed under a light cycle with lights on from 7:00 pm to 7:00 am and off from 7:00 am to 7:00 pm. Brain slices were prepared between 10:00 and 11:00 am and used for electrophysiological experiments between 2:00 and 5:00 pm, corresponding to ZT 15-16 and ZT 19–22, respectively.

We performed electrophysiological experiments like those conducted for daytime SCN neurons. Consistent with the results observed in daytime neurons, nighttime SCN neurons showed no differences in *Rm*, RMP, or the frequency of APs induced by current injection between knockout mice and their littermate controls (Fig. 5a, b). However, unlike daytime neurons, nighttime *MT_1_^-/-^*and *Slo1^-/-^* neurons did not display significant differences in any AP-related parameters compared to their littermate controls for APs induced by both 15-pA (Fig. 5) and 20-pA (Supplementary Fig. 4) current injections, except for a significant reduction in the minimum slope in *Slo1^-/-^* neurons relative to *Slo1^+/+^*controls at 20-pA current injection.

**Fig. 5.**
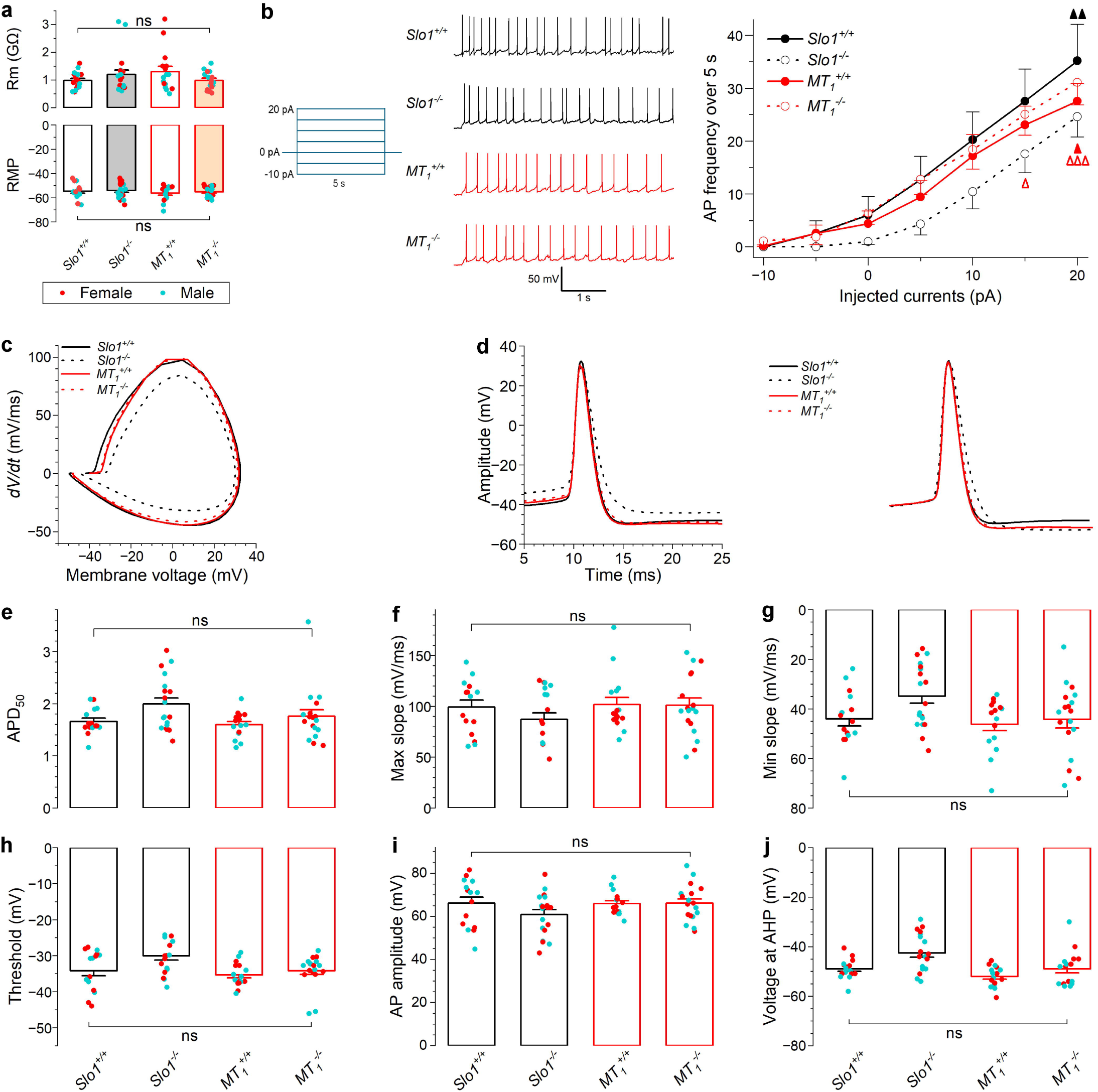
*Slo1^-/-^*or *MT_1_^-/-^* does not alter action potential (AP) duration in SCN neurons during the subjective night. **a** Membrane resistance (*R_m_*) and resting membrane potential (RMP) are unaffected by *Slo1^-/-^*or *MT_1_^-/-^.* **b** *Slo1^-/-^* or *MT_1_^-/-^* does not alter the frequency of APs induced by current injections. *Left*: Diagram of the current injection protocol. *Middle*: Representative current-clamp traces during 15-pA current injection. *Right*: AP frequency as a function of current injection. Filled and open triangles indicate statistically significant differences compared to the AP frequency at 0-pA current injection for the same group (Δ *p* < 0.05; ΔΔ *p* < 0.01; ΔΔΔ *p* < 0.001). Statistical analyses were performed using two-way mixed-design *ANOVA* followed by Tukey’s post hoc test. **c** Voltage phase plots of group averaged APs induced by 15-pA current injection. **d** Group-averaged APs induced by 15-pA current injection, displayed as actual (left) and normalized (right) waveforms. **e-j** Quantitative comparisons of AP-related parameters. Asterisks indicate statistically significant differences between groups (* *p* < 0.05; *** *p* < 0.001), while “ns” indicates no significant difference. Statistical analyses were performed using two-way *ANOVA* (factors: genotype and time of day) followed by Tukey’s post hoc test. The number of neurons recorded (*n*) were: *Slo1^+/+^* 15, *Slo1^-/-^* 19, *MT_1_^+/+^*16, and *MT_1_^-/-^* 18.

There were no apparent sex-specific differences in the results, as indicated by the distribution of male and female data points in the bar graphs (Fig. 5 and Supplementary Fig. 4). Overall, these findings suggest that Slo1 and MT_1_ do not significantly influence the AP waveform in nighttime SCN neurons, highlighting a sharp contrast to their roles in daytime SCN neurons.

We also compared AP-related parameters in SCN neurons between daytime and nighttime within each genotype. In both *Slo1^-/-^* and *MT_1_^-/-^* mice, APD50 was significantly longer during the daytime than at nighttime, whereas no difference was observed in their littermate controls (Supplementary Fig. 5). Similarly, AP minimum slope and AHP voltage were significantly smaller and less negative, respectively, during the daytime than at nighttime in *Slo1^-/-^* and *MT_1_^-/-^* mice, but remained similar between daytime and nighttime in controls (Supplementary Fig. 5). These findings indicate that the *Slo1^-/-^* and *MT_1_^-/-^*mutations introduce daytime versus nighttime differences in AP and AHP properties.

### Slo1 expression in the SCN is elevated during the daytime and reduced at nighttime

To explore the cause of the differing effects of *Slo1* and *MT_1_*knockout on AP properties in SCN neurons between the daytime and nighttime, we performed immunohistochemistry on coronal brain slices containing the SCN. Brain tissue was collected at ZT 7-8 and ZT 19–20 as daytime and nighttime samples, respectively. Immunohistochemistry was performed using an antibody against the last 100 amino acid residues of Slo1 (APC-021, Alomone Labs, Jerusalem, Israel). This epitope is present in three of the thirteen known Slo1 isoforms in the CBA/CaJ mice (<u>Gene:</u> <u>Kcnma1 (MGP_CBAJ_G0020762) - Summary - Mus_musculus_CBA_J - Ensembl</u> <u>genome browser 113</u>).

Slo1 immunoreactivity was detected in both daytime and nighttime SCN samples from *Slo1^+/+^* mice. However, the signal in the nighttime SCN was nearly 70% weaker than that in the daytime SCN (Fig. 6). In *Slo1^-/-^*mice, Slo1 immunoreactivity was also observed in the daytime SCN but was approximately 90% lower compared to daytime *Slo1^+/+^* samples (Fig. 6). Thus, the single exon in-frame deletion in *Slo1^-/-^* mice substantially reduced Slo1 expression, even though it is not expected to impact the subsequent amino acid coding sequence. This finding aligns with observations in the C57BL/6 strain, where an in-frame deletion of the corresponding exon results in the loss of both Slo1 mRNA and protein in brain tissue ^23^. The reduction in Slo1 expression may arise from various mechanisms, such as less efficient translation or reduced stability of the Slo1 mRNA or protein.

**Fig. 6.**
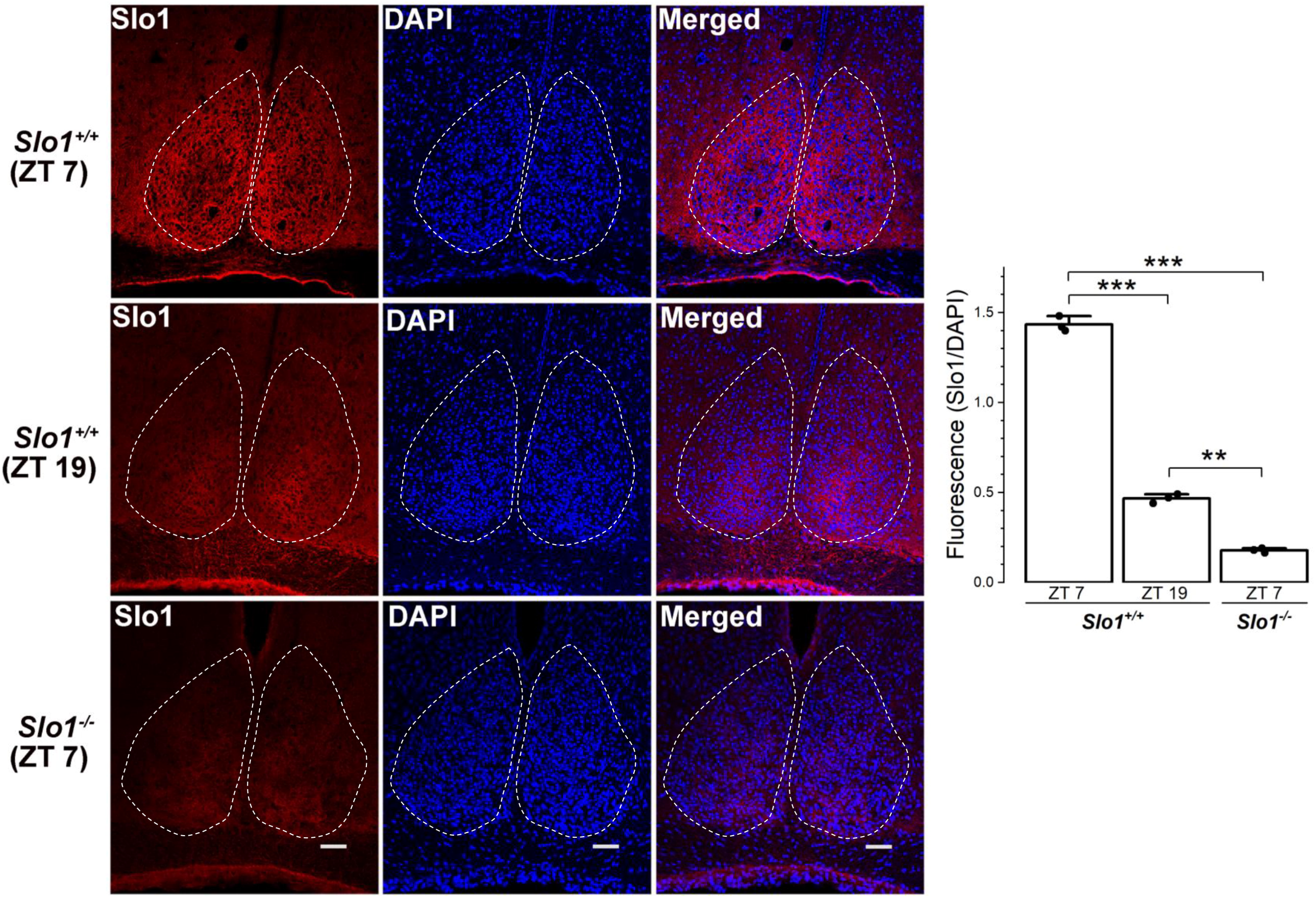
Slo1 expression in the SCN is elevated during the subjective day and reduced during the subjective night. **a** Representative immunohistochemistry images showing Slo1 immunoreactivity, DAPI staining, and merged images of the two in the bilateral SCN at ZT7 and ZT9 for *Slo1^+/+^* and at ZT7 for *Slo1^-/-^*. **b** Quantification of Slo1 immunoreactivity across groups. Asterisks indicate statistically significant differences between groups (** *p* < 0.01; *** *p* < 0.001). Statistical analyses were performed using one-way *ANOVA* followed by Tukey’s post hoc test. The sample size (*n*) was 3 animals per group, with the Slo1 immunoreactive signal (normalized to DAPI signal) for each animal derived from the average of five sections.

These results suggest that Slo1 in the SCN consists primarily or exclusively of one or more isoforms containing the C-terminal region recognized by the antibody. Moreover, Slo1 expression in the SCN is elevated during the subjective daytime and reduced during the subjective nighttime. This substantial daily variation in Slo1 expression levels may be a key factor underlying the differing effects of Slo1 knockout on the electrical properties of SCN neurons between daytime and nighttime.

### Specific domains in MT_1_ and Slo1 mediate their physical interaction

We investigated whether the functional effect of MT_1_ on Slo1 involves interactions and whether such interactions depend on specific domains by performing co-immunoprecipitation assays. These experiments were conducted with HEK293T cells transfected with MT_1_ and Slo1, which were tagged at their C-termini with HA and Flag, respectively. We immunoprecipitated the complexes using an HA antibody and immunoblotted them using a Flag antibody as well as an HA antibody.

We observed co-immunoprecipitation of full-length MT_1_ and Slo1 (Fig. 7a-c). To determine which part of Slo1 is important for its physical interaction with MT_1_, we tested whether Slo1 interacts with MT_1_ through its N- or C-terminal domain. We observed MT_1_ co-immunoprecipitation with Slo1’s N-terminal portion (amino acids 1-388), which includes all residues up to the end of S6, but not its C-terminal portion (amino acids 389-1164), which comprises the entire cytosolic domain after S6 (Fig. 7a). These results indicate that the N-terminal portion of Slo1 mediates physical interaction with MT_1_.

**Fig. 7.**
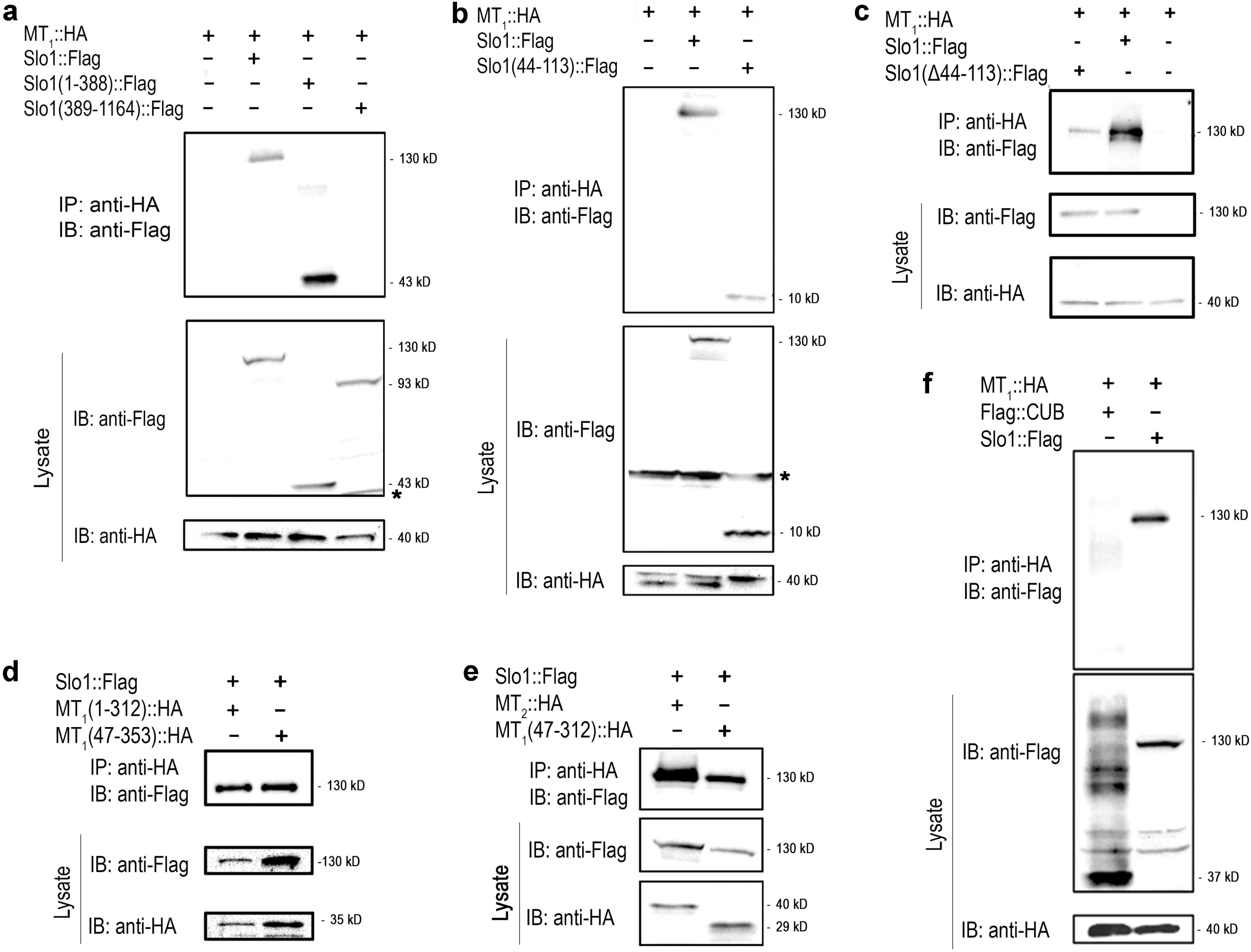
MT_1_ and Slo1 physically interact, with the membrane-spanning region of MT_1_ and the S0-S1 loop of Slo1 being critical for the interaction. HEK293T cells were transfected with HA-tagged MT_1_ (MT_1_::HA) and Flag-tagged Slo1 (Slo1::Flag) or their variants. Each gel image is representative of two independent experiments. **a** Full-length Slo1 and Slo1(1-388), but not Slo1(389-1164), coimmunoprecipitated with MT_1_. **b** Slo1(Δ44-113), which lacks a large portion of the S0-S1 loop, exhibited a much weaker interaction MT_1_ compared to full-length Slo1. **c** The Slo1(44-113) peptide coimmunoprecipitated with MT_1_. **d** Both MT_1_(Δ1-46) and MT_1_(Δ313-353), which lack the N- and C-termini, respectively, coimmunoprecipitated with Slo1. **e** MT_1_(47-312), which lacks both the N- and C-termini, as well as MT_2_, coimmunoprecipitated with Slo1. **f** An unrelated transmembrane protein, CUB3, did not coimmunoprecipitate with MT_1_.

Since the S0-S1 loop plays a key role in Slo1 regulation by several other proteins, including Slo1 β subunits ^24^, NMDA receptors ^25^, and syntaxin 1A ^26^, we tested whether the S0-S1 peptide itself (amino acids 44–113) could coimmunoprecipitate with MT_1_ and observed a positive result (Fig. 7b). We then examined whether Slo1_ΔS0-S1_, where amino acids 44-113 are deleted, could still interact with MT_1_. Slo1_ΔS0-S1_ exhibited much weaker interaction with MT_1_ compared to full-length Slo1 (Fig. 7c). These findings indicate that the S0-S1 loop is critical to Slo1 interactions with MT_1_.

To identify the domain in MT_1_ important for physical interaction with Slo1, we tested the effects of various MT_1_ deletions. We observed that two MT_1_ constructs, MT_1(1-312)_ and MT_1(47-353)_, in which the C- and N-termini were deleted, respectively, still coimmunoprecipitated with Slo1 (Fig. 7d), suggesting that the membrane-spanning region might be responsible for the physical interaction. To confirm this, we tested an MT_1_ fragment lacking both the N- and C-termini, MT_1(47-312)_. This construct also co-immunoprecipitated with Slo1 (Fig. 7e). These results suggest that the membrane-spanning region of MT_1_ mediates physical interaction with Slo1.

MT_1_ and MT_2_ are highly divergent in their N- and C-termini but highly homologous in the membrane-spanning region (Supplementary Fig. 6). The similarity in the membrane-spanning region prompted us to test whether MT_2_ can physically interact with Slo1. Indeed, we observed coimmunoprecipitation of MT_2_ with Slo1 (Fig. 7e). The finding that both MT_1_ and MT_2_ physically interact with Slo1, but only MT_1_ mediates melatonin activation of Slo1, suggests that the distinct functional effect of MT_1_ is not due to a unique ability to bind Slo1, but rather results from effects of its distinct N- or C-terminus.

Finally, we validated the specificity of our assays by testing whether MT_1_ can coimmunoprecipitate with an unrelated membrane protein. Specifically, we tested whether the CUB domain of human BAI3 (brain-specific angiogenesis inhibitor 3) with a membrane GPI anchor can coimmunoprecipitate with MT_1_. We were unable to detect a CUB signal in the immunoprecipitates obtained with MT_1_ (Fig. 7f). This result confirms that the interactions observed between MT_1_ and Slo1 are specific.

## DISCUSSION

Our study defines a circadian mechanism in which melatonin promotes sleep via MT_1_-dependent activation of Slo1 in SCN neurons. Key findings include: (1) Ablation of Slo1 and MT_1_ reduces sleep duration and prolongs APs in SCN neurons, with both effects occurring exclusively or primarily during the subjective day; (2) Slo1 expression peaks during the day, coinciding with its sleep-promoting role; (3) MT_1_-Slo1 interaction requires specific structural domains; and (4) global Slo1 deletion causes seizures, underscoring its neurophysiological roles beyond sleep regulation. Our results thus define new roles for MT_1_ and Slo1 in sleep.

EEG and EMG recordings demonstrated that both *Slo1^-/-^*and *MT_1_^-/-^* mice exhibited reduced REM and NREM sleep, with effects occurring during the subjective day. This aligns with our finding that Slo1 expression is highest during the day, and with prior work showing daytime peaks in MT_1_ expression in rodent brains ^27^. Thus, Slo1 and MT_1_ appear to regulate sleep during the resting phase of the circadian cycle.

The observed effects of *MT_1_^-/-^* and *Slo1^-/-^*on SCN neuronal properties suggest the presence of either constitutively active MT_1_ or locally synthesized melatonin in brain slices under our experimental conditions. As for the first possibility, studies involving inverse receptor agonists have revealed that a variety of GPCRs, including MT_1_, can exhibit constitutive activity in the absence of ligands ^28–30^. Although MT_1_ does not alter Slo1 activity in the absence of melatonin in *Xenopus* oocytes ^17^, this finding does not rule out the possibility that MT_1_ may be constitutively active in native SCN neurons. As for the second possibility, any melatonin involved in MT_1_ activation in the brain slice for electrophysiological recording must have a local source, as it is unlikely that melatonin from the pineal gland would remain after the >2-hour incubation in artificial cerebrospinal fluid prior to the recording. Previous studies suggest that melatonin can be synthesized locally in brain tissues. For example, mRNAs encoding the key enzymes for melatonin synthesis, arylalkylamine *N*-acetyltransferase (AANAT) and hydroxyindole-*O*-methyltransferase (HIOMT), are detected in various rat brain regions ^31,32^. Melatonin is detected in brain tissues of pinealectomized rats ^27^, with the highest levels in the hypothalamus ^33^. Furthermore, there is evidence that astrocytes, glial cells, and neural stem cells can synthesize melatonin ^34–36^. Since the *K_i_* of melatonin for MT_1_ is 0.1 nM ^1,37^ and the *EC_50_* for melatonin’s activation of Slo1 via MT_1_ in the *Xenopus* oocyte heterologous expression system is 0.77 nM ^17^, even sub-nanomolar concentrations of melatonin in the SCN brain slice is sufficient to activate Slo1 via MT_1_.

Deletion of *Slo1* or *MT_1_* did not significantly alter AP frequencies in SCN neurons, indicating that these genes regulate sleep through mechanisms independent of modulation of AP frequency modulation. This contrasts with earlier studies reporting increased nighttime SCN firing rates in *Slo1^-/-^*mice ^5,38^. However, those studies used multi-electrode arrays ^38^ or single-unit recordings ^5^, methodologies that are not directly comparable to our whole-cell approach, which recorded APs evoked by controlled current injections. Given the functional heterogeneity of SCN neurons, the subpopulations exhibiting increased AP firing rates in prior multi-electrode array experiments ^38^ likely differ from those analyzed here. Furthermore, the single-unit method required extensive neuronal sampling to detect significant differences in spike rates due to high variability in neuronal activity ^5^. These methodological and cellular distinctions likely explain why we observed no effect of Slo1 or MT_1_ ablation on AP frequencies.

Consistent with the findings of epilepsy in human patients with loss-of-function mutations in *Slo1* ^21,22^, we observed robust seizure activity in *Slo1^-/-^* mice. Our findings align with a previous study reporting seizure-like locomotor behavior in *Slo1^-/-^* mice ^22^, but contrast with another study reporting no difference in pentylenetetrazol-induced seizures between wild-type and *Slo1^-/-^*mice ^39^. Direct comparison of our results with those of the previous studies is challenging because they used mice strains that either do not secrete melatonin or with an unknown capacity for melatonin synthesis. The discovery of a strong association between *Slo1^-/-^* and seizures in the CBA/CaJ strain suggests that this strain can be particularly valuable for investigating the cellular and molecular mechanisms underlying seizures associated with Slo1 loss-of-function mutations. In summary, our findings suggest that melatonin promotes sleep by enhancing AP repolarization in SCN neurons through MT_1_-mediated activation of Slo1. This effect is facilitated by a physical interaction between MT_1_ and Slo1. A key question is how the modulation of AP waveforms by MT_1_ and Slo1 in SCN neurons translates into a sleep-promoting effect. In *C. elegans*, a specific melatonin receptor and SLO-1 function together to downregulate neurotransmitter release to promote the DTQ ^17^. Conceivably, MT_1_ and Slo1 might also function together to regulate neurotransmitter release from SCN neurons to promote sleep. Future studies are needed to elucidate the precise neural circuit and synaptic mechanisms by which MT_1_ and Slo1 in the SCN enhance sleep.

## Acknowledgements

This work was supported by NIH grants R01MH085927 and R01NS109388 (both to Z.W.W.). The authors express sincere gratitude to Yinyying Ge for her guidance on EEG/EMG electrode installation, Bojun Chen for his guidance on coimmunoprecipitation assays, David Martinelli for providing the FLAG-tagged CUB plasmid, and Katie Lowther for her guidance on genotyping.

## Author contributions

Z.W.W. contributed to the conception, design, interpretation of results, and manuscript writing. K.V. contributed to genotyping, experimentation, and data analysis for Figures 1-3, 6, and 7, as well as manuscript writing. A.A. performed the experimentation and data analysis for Figures 4, 5, and S2–S5. L.N. contributed to the initial preliminary studies and provided guidance and training to A.A. in experimentation. Y.S. contributed to the initial preliminary studies and assisted in training K.V. in molecular biological techniques. F.L.C. helped with the adoption of the slice recording technique in Z.W.W.’s lab and provided troubleshooting support. X.M.M. worked together with K. V. in the immunohistochemistry experiments. D.K. and S.P.Y. assisted in maintaining and genotyping mouse strains. X.H. guided K.V. in EEG/EMG recordings and analyses.

## METHODS

### Creation of *MT_1_* and *Slo1* knockout mice

*Mtnr1a (MT_1_)* and *Kcnma1 (Slo1)* knockout mice were generated by the Jackson Laboratory (JAX) using CRISPR-mediated gene editing of zygotes derived from CBA/CaJ mice (JAX Stock# 000654). Briefly, the Benchling CRISPR Guide RNA Design Tool (https://www.benchling.com/crispr) was used to identify CRISPR target sites for excising exon 1 of *MT_1_* and exon 8 of *Slo1* with minimum off-target potential. The following CRISPR guide RNA (crRNA) sequences were identified:

For *MT_1_*: *MT_1__*crRNA1 (5’-GGG CCA TTG TCC TGT CGC CC) and *MT_1_*_crRNA2 (5’-CTC CTG GAT GAT CTT TGT CA).

For *Slo1*: *Slo1_*crRNA1 (5’-ATC AAG CAT CAG CGA TGC GT) and *Slo1_*crRNA2 (5’-GGC ACC TGC CCC TAA AGA GC).

Single guide RNAs (sgRNAs) with sequences specific to *MT_1_* and *Slo1* were purchased from Synthego (Redwood City, CA, USA), and SpCas9 protein was obtained from Integrated DNA Technologies (Coralville, IA, USA). These components were electroporated into zygotes following an established method.^40^

Live-born pups were initially screened by PCR using the following primer pairs:

- For *MT_1_*: *MT_1_*_F1 (5’-CTT GAT GCC TCC ACG TGT CT) and *MT_1_*_R1 (5’-TTT TTG GCG GCT TCT GCA TC), which amplify an 827 bp fragment from the wild-type allele and a ∼400 bp fragment from the knockout allele.
- For *Slo1*: *Slo1*_F1 (5’-CAA GCA GTC CCA GAA GCA GA) and *Slo1*_R1 (5’-TGA AGC AAC TGT TCC CTG GA), which amplify a 1207 bp fragment from the wild-type allele and a ∼400 bp fragment from the knockout allele.

Potential founders were further confirmed by PCR followed by Sanger sequencing of the PCR products using the primers described above. Confirmed founders, including an *MT_1_* allele with a 500 bp deletion and a *Slo1* allele with an 866 bp deletion and 3 bp insertion, were bred with CBA/CaJ mice to establish stable knockout lines for further characterization. The sequences, annealing sites, and PCR product sizes of oligo pairs for routine PCR genotyping are shown in Fig. 1.

Mice were housed in a controlled environment with a 12-hour light/dark cycle and provided with a standard diet and water ad libitum. All experiments were conducted in accordance with the National Institutes of Health (NIH) guidelines and were approved by the Institutional Animal Care and Use Committee (IACUC) at UConn Health.

### Mouse stereotaxic surgery and EEG/EMG recordings

A 12- to 14-week-old mouse was anesthetized using 2% isoflurane delivered via a gas evacuation apparatus (RWD, Gas Evacuation Apparatus R546W). The anesthetized mouse was then positioned in a stereotaxic frame equipped with a mouse adaptor (51509M, Stoelting Co., Wood Dale, IL, USA). Following exposure of the skull, an EEG/EMG head mount featuring two EEG inputs and one EMG input (8201-SS, Pinnacle Technology, Parsippany, NJ, USA) was affixed to the skull using four stainless steel screws (two anterior screws: 8209; two posterior screws: 8212; Pinnacle Technology, Parsippany, NJ, USA). The exposed skull and neck regions were subsequently covered with a ceramic dental compound (51458, Stoelting Co., Wood Dale, IL, USA).

After surgery, the mouse was allowed to recover for 7–10 days before being transferred to an EEG/EMG recording chamber. The chamber was maintained on a 12-hour light/dark cycle (lights on from 7:00 AM to 7:00 PM). The head mount was connected to a Data Acquisition and Control System (8206, Pinnacle Technology) via a preamplifier (8202-SE, Pinnacle Technology). Continuous EEG and EMG signals, along with synchronized video recordings, were acquired for 72 hours without human intervention. Data were amplified and recorded using Sirenia Acquisition software (Pinnacle Technology).

### Sleep and wake duration quantification

Sleep and wake durations were quantified using the final 24 hours of EEG and EMG recordings. EEG traces were automatically segmented into 4-second epochs and classified as wake, non-rapid eye movement (NREM) sleep, or rapid eye movement (REM) sleep using a cluster scoring approach implemented in Sirenia Sleep Pro software (Pinnacle Technology). The automated classifications were manually verified by cross-referencing raw EEG/EMG data and corresponding spectral plots (power over frequency).

Wake periods were characterized by high-amplitude EMG activity and a dominant EEG frequency greater than 4 Hz. NREM sleep was identified by a dominant EEG frequency below 4 Hz and low-amplitude EMG activity. REM sleep was distinguished by a dominant EEG frequency between 5 and 8 Hz accompanied by low-amplitude EMG signals. The total time spent in wake, NREM, and REM states during both the light and dark phases was calculated and extracted using the software.

### Seizure identification and quantification

Ictal events were identified and quantified from the final 24 hours of EEG/EMG recordings in *Slo1*^⁻/⁻^ mice using Sirenia Seizure Pro software (Pinnacle Technology). Ictal discharges were defined as rhythmic bursts of sharply contoured EEG activity lasting at least 10 seconds. Each event was manually selected within the EEG trace, and the total power associated with the event was quantified using Sirenia Seizure Pro. To confirm the behavioral state during each event, EMG signals and synchronized video recordings were reviewed to ensure that the mouse was awake at the time of the seizure.

### Brain slice electrophysiology

Mice aged 3–5 weeks were anesthetized with 2% isoflurane and decapitated at approximately 11:00 AM. The brain was rapidly removed from the cranium and placed in an ice-cold tray containing carbogenated slicing solution (composition in mM: 125 NaCl, 2.5 KCl, 1.25 NaH₂PO₄, 26 NaHCO₃, 10 glucose, 10 sucrose, 0.5 CaCl₂·2H₂O, and 2 MgCl₂·6H₂O). The brain was then mounted on a specimen holder (VF-SPS-VM-12.5-BOS) and embedded in 2% melted agarose (A9414, Sigma) maintained at 42°C using an Eppendorf ThermoMixer F2.0. Coronal slices of 300 µm thickness were cut using an ENDURIUM® blade (EF-INZ10, Cadence, San Jose, CA, USA) on a *Compresstome* (VT-510-0Z, Precisionary Instruments LLC, Ashland, MA, USA) in ice-cold slicing solution. The entire procedure, from decapitation to obtaining brain slices, was completed within approximately 15 minutes. Slices containing the suprachiasmatic nucleus (SCN) were incubated at 37°C for 30 minutes and then maintained at room temperature in carbogenated maintenance solution (composition in mM: 125 NaCl, 2.5 KCl, 1.25 NaH₂PO₄, 26 NaHCO₃, 10 glucose, 10 sucrose, 2 CaCl₂·2H₂O, and 4 MgSO₄·7H₂O) until used for electrophysiological recordings.

SCN-containing slices were transferred to a recording chamber (RC-26G, Warner Instruments, Hamden, CT, USA) mounted on a magnetic platform (PM-1, Warner Instruments). The chamber was continuously perfused with carbogenated artificial cerebrospinal fluid (aCSF; composition in mM: 125 NaCl, 2.5 KCl, 1.25 NaH₂PO₄, 26 NaHCO₃, 10 glucose, 10 sucrose, 2 CaCl₂·2H₂O, and 1.3 MgCl₂·6H₂O), with osmolarity adjusted to 305 mOsm using sucrose. The intracellular pipette solution contained (in mM): 125 K-gluconate, 4 KCl, 0.2 CaCl₂·2H₂O, 10 HEPES, 1 EGTA, 10 phosphocreatine-Na₂, 4 Na₂-ATP, and 0.3 Na-GTP, with osmolarity adjusted to 290 mOsm using sucrose.

SCN neurons were visualized using an infrared monochrome video camera (IR-2000, DAGE-MTI, Michigan City, IN, USA) under IR-DIC illumination, controlled by IR-Capture software (DAGE-MTI) on an upright microscope (Axio Examiner.Z1, Zeiss, Germany). Tear-drop or balloon-shaped neurons in the ventromedial SCN were selected for whole-cell patch-clamp recordings. Membrane resistance and capacitance were measured using the Membrane Test function, and action potentials (APs) were recorded in response to current injections. Neurons with an access resistance of <25 MΩ were selected for recording. Recordings were included for analysis only if the access resistance remained stable, increasing by no more than 10% during the experiment. Electrophysiological recordings were performed using a Multiclamp 700B amplifier, a digital-to-analog converter (Digidata 1550B), and pCLAMP software (v11, Molecular Devices, Sunnyvale, CA, USA). Data were low-pass filtered at 2 kHz and acquired at 10 kHz. Recording electrodes (tip resistance 4–6 MΩ) were pulled from borosilicate glass capillaries (BF120-69-10, Sutter Instrument, Novato, CA, USA).

Raw AP data files acquired using Clampex were imported into *Igor Pro 9* (WaveMetrics) for analysis. For each current injection step, the software automatically quantified key parameters for individual APs, including APD₅₀, maximum/minimum slopes, threshold, amplitude, AHP voltage, and the timing of AP peaks and AHP occurrences. Results were exported to *Excel* to calculate average values for all APs and tally the number of APs per current injection step.

AHP voltage for *MT_1_^+/+^* and *Slo1^+/+^*mice was automatically quantified by *Igor Pro 9*, while that for *MT_1_^-/-^* and *Slo1^-/-^* mice was estimated using the following approach. First, the time interval between the AP peak and the AHP voltage was calculated from the automatically detected times for AP peak and AHP for *MT_1_*⁺*/*⁺ and *Slo1*⁺*/*⁺ mice in *Igor Pro 9*, yielding a interval of ∼5 ms. For *MT*^1⁻/⁻^ and *Slo1*^⁻/⁻^ mice, AHP voltage was defined as the membrane voltage measured 5 ms post-AP peak in averaged AP traces (per current injection step), identified using ClampFit (Molecular Devices).

### Immunohistochemistry

Mice were anesthetized with 2% vaporized isoflurane and subjected to transcardial perfusion. Perfusion began with phosphate-buffered saline (PBS) for 5 minutes, followed by 4% paraformaldehyde (PFA) until the tail stiffened and the liver turned pale, indicating proper fixation. The brain was then removed and post-fixed in ice-cold 4% PFA for 2 hours. After fixation, the brain was cryoprotected by incubation in 20% sucrose solution at 4°C overnight. Coronal sections of 14 µm thickness were cut using a Leica CM1860 UV cryostat (Leica, USA) and mounted onto gelatin-coated Superfrost microscope slides (12-550-123, Fisher Scientific, USA).

Slides were washed three times in PBS for 5 minutes each to remove residual optimal cutting temperature (OCT) compound. Sections were then blocked for 1 hour at room temperature in PBS containing 5% goat serum, 1% bovine serum albumin (BSA), and 0.2% Triton X-100 (pH 7.4). After blocking, sections were incubated overnight at 4°C with a 1:500 dilution of rabbit polyclonal anti-BK channel antibody (APC-021, Alomone Labs, Israel). The following day, sections were washed three times in PBS for 5 minutes each and incubated for 1 hour at room temperature with Alexa Fluor 594-conjugated secondary antibody (Thermo Fisher Scientific, USA). After secondary antibody incubation, sections were washed three more times in PBS for 5 minutes each and mounted with DAPI Fluoromount-G (Southern Biotech, 0100-20, USA). Images were acquired using a Zeiss LSM 800 confocal microscope (Zeiss, Germany).

For quantification, images were imported into ImageJ software. The scale was set using metadata exported from ZEN 2.7 software (Zeiss, Germany). The bilateral suprachiasmatic nuclei (SCN) were manually selected for analysis. The integrated density of the red fluorescence signal (Slo1) and the DAPI signal (nuclei) were measured. The integrated density of the Slo1 signal was normalized to the corresponding DAPI signal. Normalized values were exported to Origin software (OriginLab, USA) for calculating the mean ± standard error (SE) and generating graphical representations.

### Plasmids

The plasmids encoding hemagglutinin (HA)-tagged mouse MT_1_ (*wp2117*) and FLAG-tagged mouse Slo1 (*wp2116*) were constructed by cloning PCR-amplified products into the pIRES2– EGFP vector (Clontech, Mountain View, CA, USA). The *Mtnr1a* cDNA (NM_008639, MR224599, OriGene, Rockville, MD, USA) and *Slo1* cDNA (MBr5/3, provided by Dr. Lawrence Salkoff’s lab) were amplified using primer pairs 4378/4379 and 4376/4377, respectively (Table 1). The antisense primers were designed to fuse the 3’-ends of the MT_1_ and Slo1 coding sequences in-frame with the HA and FLAG epitope tags, respectively. The resulting PCR products were cloned into the pIRES2–EGFP vector at BglII/PstI (MT_1_) and KpnI/BamHI (Slo1) restriction sites. To generate plasmids encoding HA-tagged MT_1_(1-312) (*wp2144*) and FLAG-tagged Slo1(1-388) (*wp2138*), the corresponding cDNA fragments were amplified using primer pairs 4378/4433 and 4397/4419, respectively (Table 1). The antisense primers 4433 and 4419 were designed to fuse the codons for amino acids 312 (MT_1_) and 388 (Slo1) in-frame with the HA and FLAG tags, respectively. The amplified products were cloned into the pIRES2-EGFP vector at BglII/PstI (MT_1_) and SalI/BamHI (Slo1) restriction sites.

Plasmids encoding HA-tagged MT_1_(47-353) (*wp2143*) and FLAG-tagged Slo1(389-1164) (*wp2139)* were generated by amplifying the corresponding cDNA fragments using primer pairs 4432/4379 and 4420/4377, respectively (Table 1). The sense primers 4432 and 4420 included start codons (ATG). The amplified products were cloned into the pIRES2-EGFP vector at BglII/PstI (MT_1_) and SalI/BamHI (Slo1) restriction sites. The plasmid encoding HA-tagged MT_1_(47-312) (*wp2151)* was generated by amplifying the corresponding cDNA fragment using primer pair 4432/4433, where the sense primer 4432 included a start codon (ATG) and the antisense primer 4433 introduced a premature stop codon. The amplified product was cloned into the pIRES2-EGFP vector at BglII/PstI restriction sites.

The plasmid encoding FLAG-tagged Slo1(44-113) (*wp2152)* was generated by amplifying the *Slo1* cDNA using primer pair 4478/4479, with a methionine codon added before the codon for amino acid 44. The antisense primer fused the codon for amino acid 113 in-frame with the FLAG tag. The PCR product was cloned into the pIRES2–EGFP vector at BglII/BamHI sites. The FLAG-tagged Slo1(Δ44-113) plasmid was constructed using the Hi-Fi DNA Assembly kit (E2621S, New England Biolabs, Ipswich, MA, USA). Two PCR products, amplified using primer pairs 4766/4668 and 4667/4746, were assembled with SalI/HindIII-digested plasmid (*wp2116*). The antisense primer 4668 and the sense primer 4667 were complementary to the *Slo1* cDNA sequence but excluded the coding nucleotides for amino acids 44-113.

The plasmid encoding FLAG-tagged CUB (*wp2118)* was a gift from Dr. David Martinelli ^41^.

### Coimmunoprecipitation

Human embryonic kidney (HEK) 293T cells were cultured in Dulbecco’s Modified Eagle Medium (DMEM) supplemented with 10% fetal bovine serum (FBS). Transient transfection with hemagglutinin (HA)-tagged mouse MTNR1A and FLAG-tagged mouse Slo1 (MBr5/3) and their various deletion constructs were carried out was performed using Lipofectamine™ 2000 transfection reagent (Invitrogen) according to the manufacturer’s instructions. Cells were harvested 48 hours post-transfection and lysed in 1% CHAPS buffer (75621-03-03, Fisher BioReagents) supplemented with a protease inhibitor cocktail (Roche, Basel, Switzerland), with the pH adjusted to 6.8. Lysates were centrifuged at 16,000 rpm for 15 minutes to remove cellular debris.

Supernatants were incubated with Pierce™ anti-HA magnetic beads (Thermo Fisher Scientific) for 30 minutes at room temperature with gentle rotation. The magnetic beads were collected using a Magna GrIP rack (20-400, Millipore, Jaffrey, NH, USA) and washed twice with 1X TBST buffer (20 mM Tris, pH 7.6, 150 mM NaCl, 0.1% Tween-20) and once with ultrapure distilled water (10977015, Invitrogen, Carlsbad, CA, USA). Immunoprecipitated proteins were eluted from the beads by incubation with 1X SDS loading buffer for 10 minutes at 37°C. The eluted proteins were then resolved by SDS/PAGE and analyzed by immunoblotting

### Western blot

Immunoprecipitated proteins were resolved by electrophoresis on an SDS-PAGE gel (Invitrogen, USA) and transferred onto a 0.22 µm PVDF membrane (Millipore, USA) under ice-cold conditions at 200 mA for 3 hours. Following transfer, the membrane was blocked with 5% non-fat milk in 1X TBST buffer (20 mM Tris, pH 7.6, 150 mM NaCl, 0.1% Tween-20) for 30 minutes at room temperature. The membrane was then incubated with the appropriate primary antibody overnight at 4°C.

The next day, the membrane was washed three times with TBST buffer for 5 minutes each and incubated with the corresponding horseradish peroxidase (HRP)-conjugated secondary antibody at room temperature for 1 hour. Protein signals were visualized using a ChemiDoc Imaging System (Bio-Rad, Hercules, CA).

### Antibodies and Reagents

The following primary and secondary antibodies were used for immunoblotting:

- Rabbit monoclonal anti-FLAG antibody (SAB4301135, Sigma) at a dilution of 1:2000.
- Mouse monoclonal anti-HA antibody (sc-7392, Santa Cruz Biotechnology, Dallas, TX, USA) at a dilution of 1:1000.
- Horseradish peroxidase (HRP)-conjugated goat anti-rabbit IgG (sc-2004, Santa Cruz Biotechnology) at a dilution of 1:10,000.
- HRP-conjugated goat anti-mouse IgG (sc-2005, Santa Cruz Biotechnology) at a dilution of 1:10,000.

### Statistical analysis

Experimental results are expressed as mean ± standard error (SE). Data graphing and statistical analyses were performed using OriginPro software (OriginLab, Northampton, MA, USA). Specific statistical methods used for each analysis are detailed in the corresponding figure legends.

