## Supplemental Figures for "Circadian Control of Sleep by Melatonin via MT_1_-Dependent Activation of BK Channels in the Suprachiasmatic Nucleus"

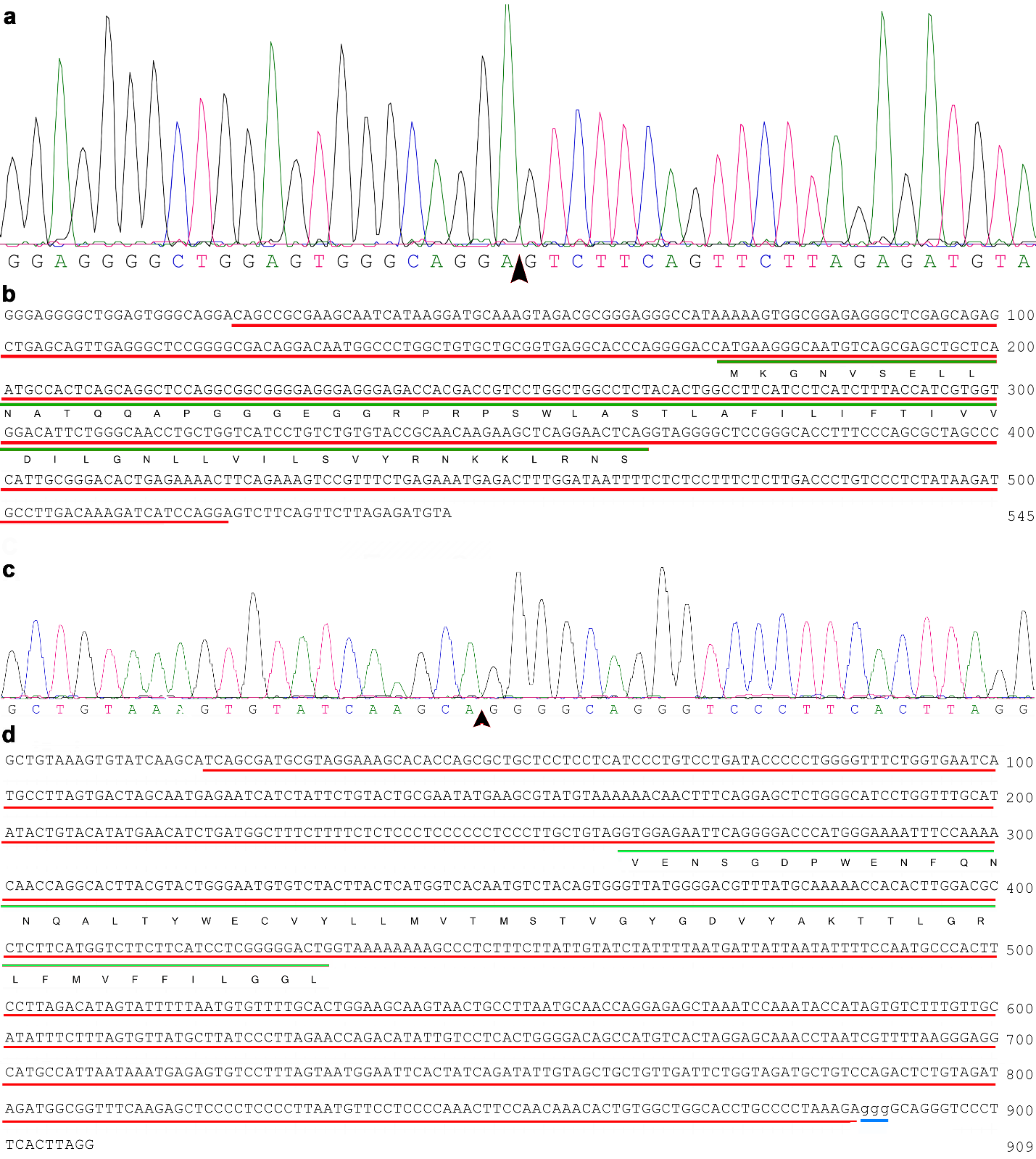

**Supplementary Fig. 1. Deletions in *MT_1_^-/-^* and *Slo1^-/-^* mice revealed by DNA sequencing. a,** Chromatogram showing the location (indicated by the arrowhead) of a 500-bp deletion in the *Mtnr1a* gene. **b,** Nucleotide sequence encompassing the 500-bp deletion (underlined in red) and the flanking nucleotides matching the chromatogram. Amino acids encoded by exon 1 are underlined in green. **c,** Chromatogram showing the location (indicated by the arrowhead) of an 866-bp deletion and a 3-bp insertion in the *Kcnma1* gene. **d,** Nucleotide sequence encompassing the 866-bp deletion (underlined in red), the 3-bp insertion (underlined in blue), and the flanking nucleotides matching the chromatogram. Amino acids encoded by exon 8 are underlined in green.

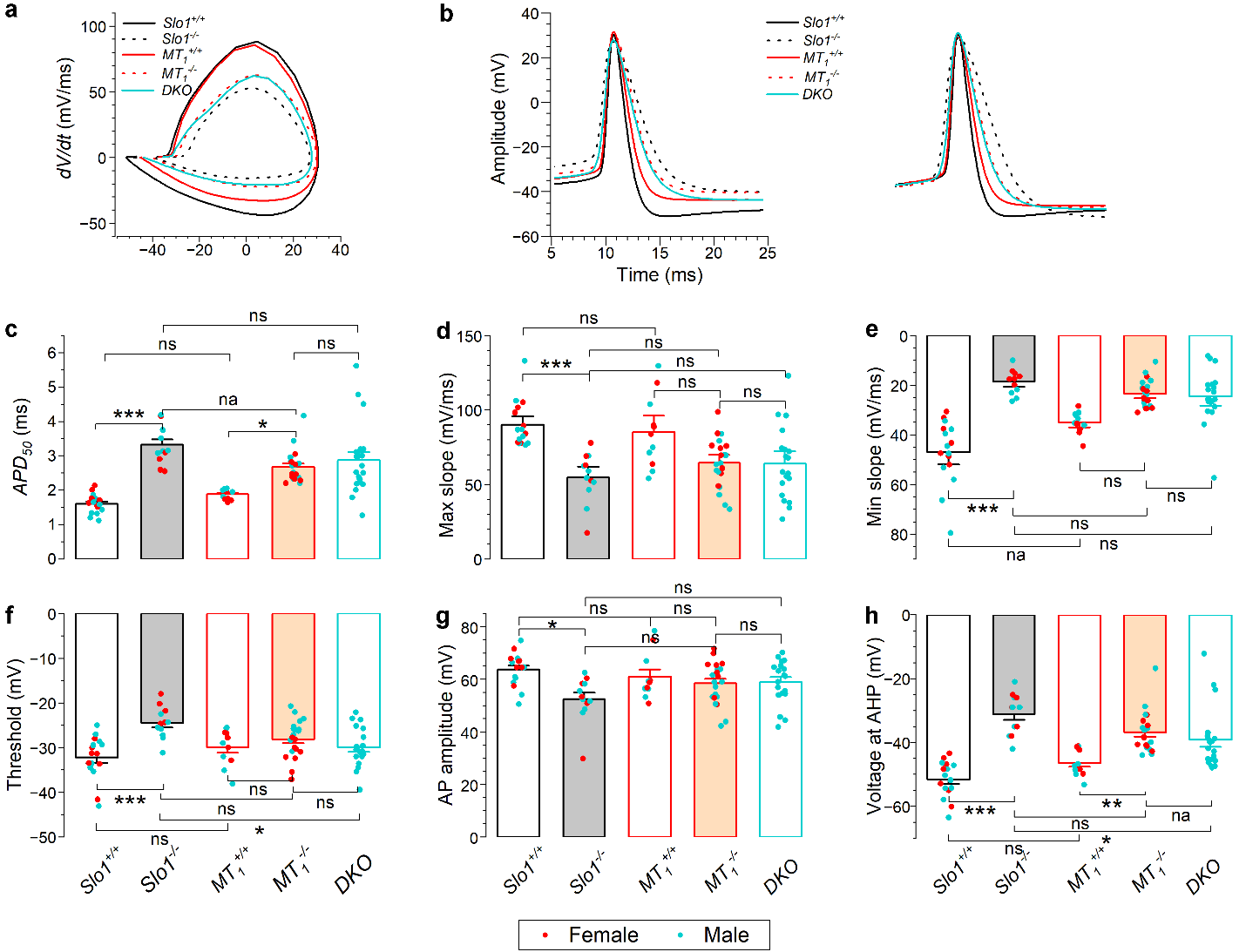

**Supplementary Fig. 2. SCN neurons in *Slo1^-/-^* and *MT_1_^-/-^* mice exhibit significant differences in APD_50_ and AHP during the subjective day.** Quantitative analysis was performed on APs induced by 20 pA current injection. **a,** Voltage phase plots of averaged APs. **b,** Averaged APs, displayed as actual (left) and normalized (right) waveforms. **c-h,** Quantitative comparisons of AP-related parameters. Asterisks indicate statistically significant differences between groups (* *p* < 0.05; ** *p* < 0.01; *** *p* < 0.001), while “ns” indicates no significant difference. Statistical analyses were performed using two-way mixed-design *ANOVA* (factors: genotype and time of day) followed by Tukey’s post hoc test. Blue and red dots represent data from male and female mice, respectively. Data were from the same experiments as in Fig. 4.

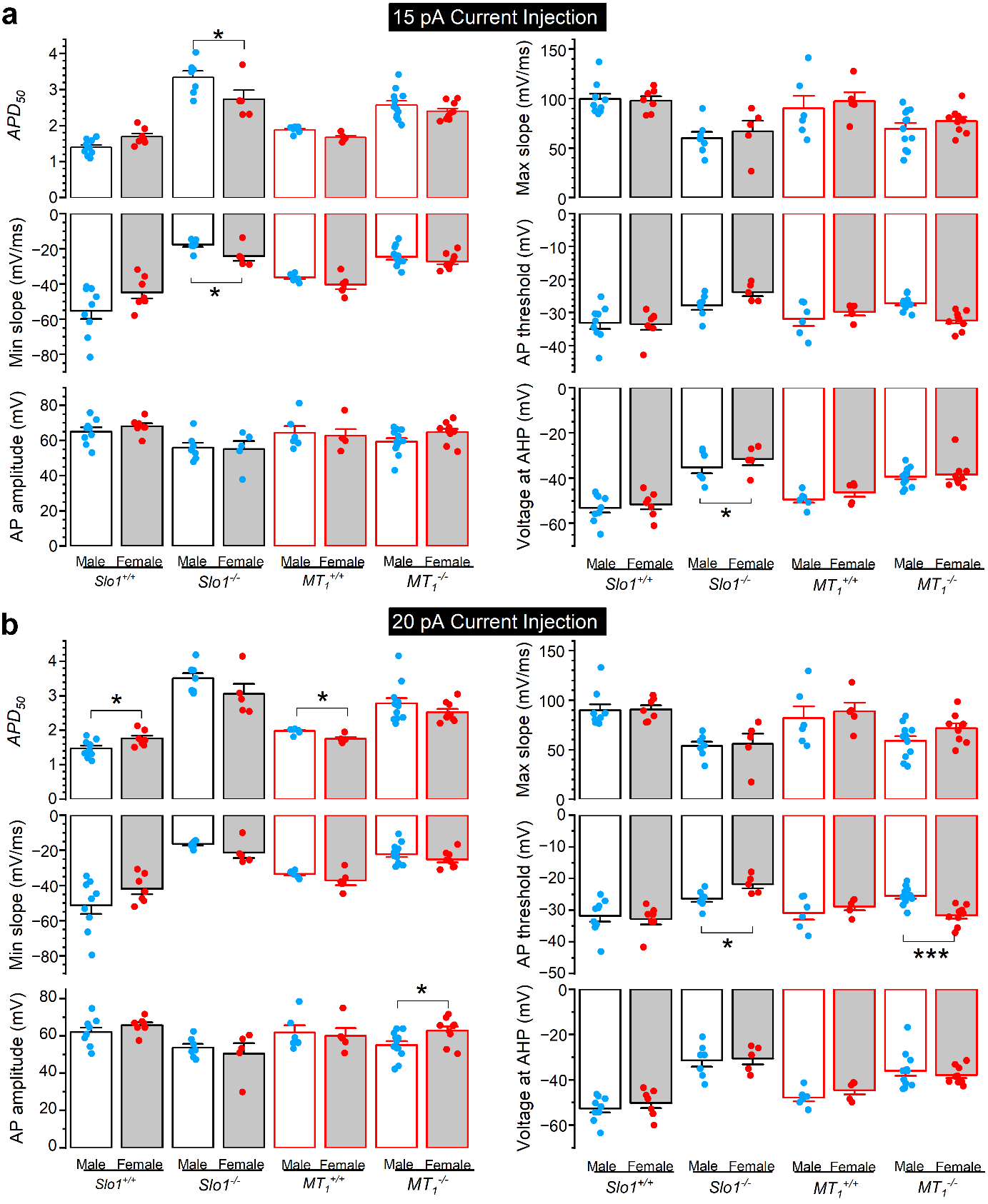

**Supplementary Fig. 3. SCN neurons in *Slo1^-/-^* and *MT_1_^-/-^* mice do not display sex-specific differences in AP-related parameters during the subjective day.** Quantitative comparisons of APs induced by 15-pA (**a**) and 20-pA (**b**) current injections between male and female mice. Statistical analyses were performed using two-way mixed-design *ANOVA* (factors: genotype and sex) followed by Tukey’s post hoc test. The data are the same as those in Fig. 4 and Supplementary Fig. 2.

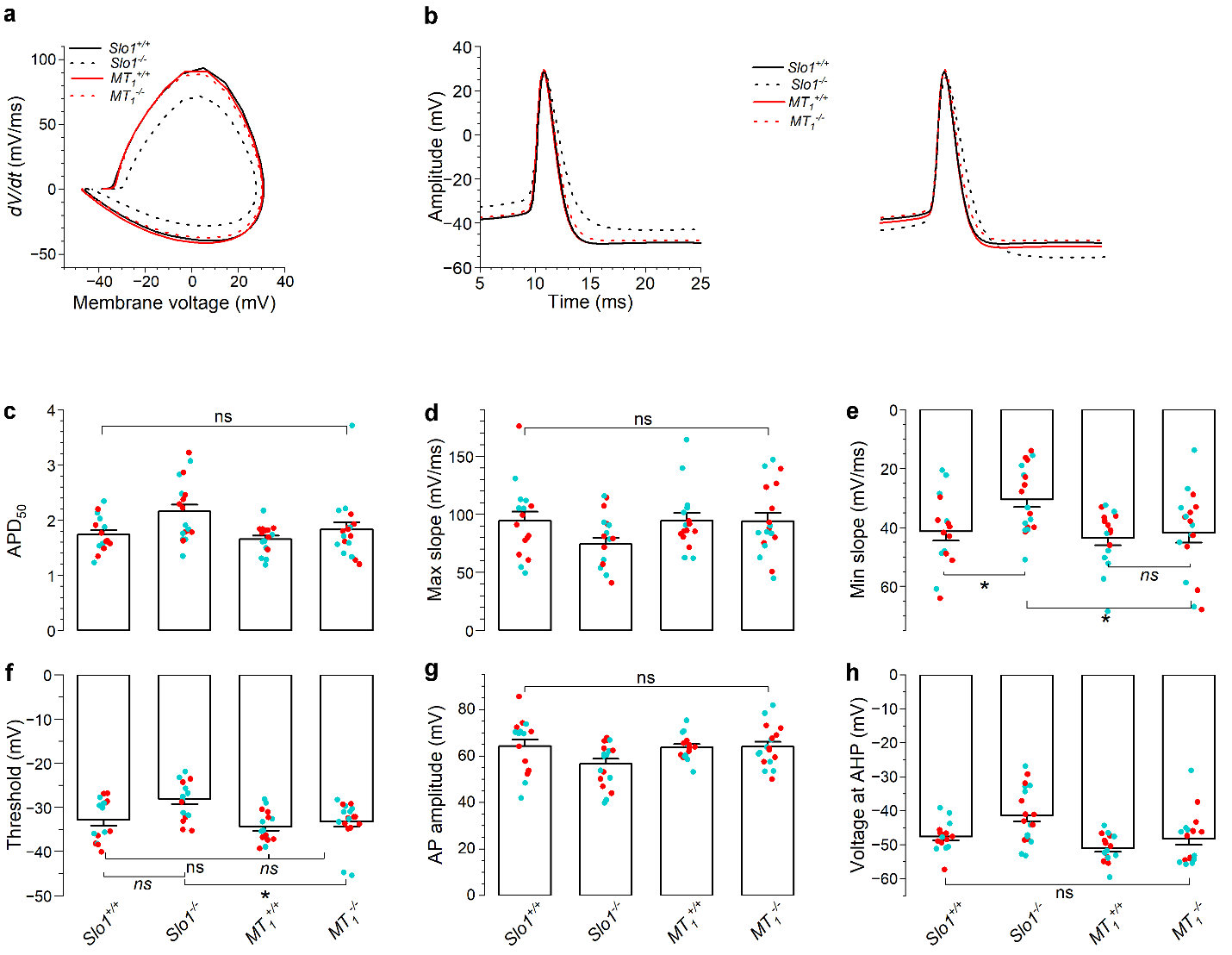

**Supplementary Fig. 4. *Slo1^-/-^* or *MT_1_^-/-^* does not alter APD_50_ and AHP during the subjective night.** Quantitative analysis was performed on APs induced by 20 pA current injection. **a,** Voltage phase plots of averaged APs. **b,** Averaged APs, displayed as actual (left) and normalized (right) waveforms. **c-h,** Quantitative comparisons of AP-related parameters. Asterisks indicate statistically significant differences between groups (* *p* < 0.05; ** *p* < 0.01; *** *p* < 0.001), while “ns” indicates no significant difference. Statistical analyses were performed using two-way mixed-design *ANOVA* (factors: genotype and time of day) followed by Tukey’s post hoc test. Blue and red dots represent data from male and female mice, respectively. The data are the same as those in Fig. 5.

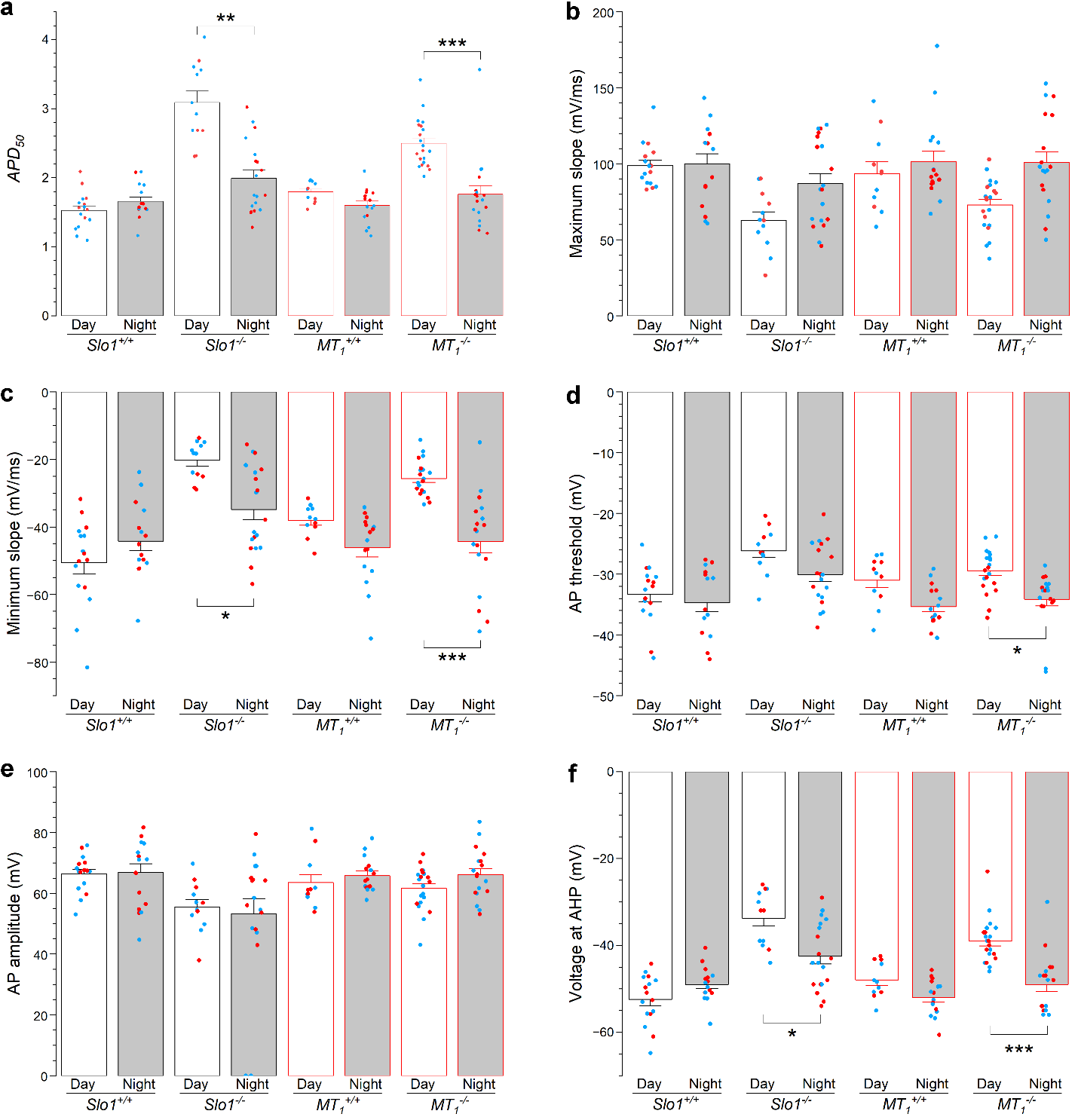

**Supplementary Fig. 5. APD_50_ and AHP display significant differences between subjective and subjective night in *Slo1^-/-^* and *MT_1_^-/-^* mice but not in their littermate controls. a-f,** Quantitative comparisons of AP-related parameters. Asterisks indicate statistically significant differences between the indicated groups (* *p* < 0.05; ** *p* < 0.01; *** *p* < 0.001), while “ns” indicates no significant difference. Statistical analyses were performed using two-way mixed-design *ANOVA* (factors: genotype and time of day) followed by Tukey’s post hoc test. The data are the same as those in Figs. 4 and 5. Blue and red dots in the graphs represent data from male and female mice, respectively.

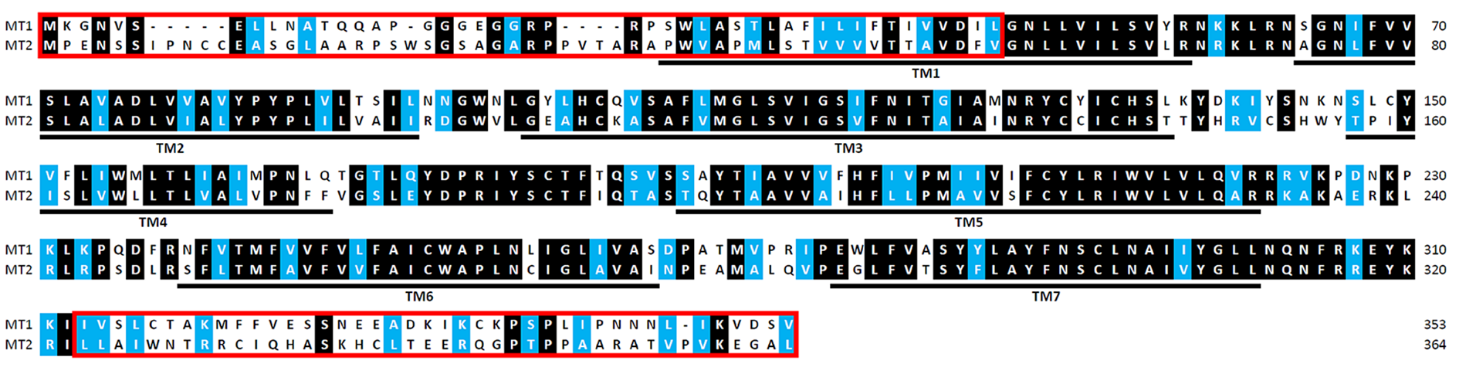

**Supplementary Fig. 6. Amino acid sequence alignment of MT_1_ and MT_2_ melatonin receptors.** Identical and similar residues are highlighted with black and blue backgrounds, respectively. Predicted transmembrane domains (TM1–TM7) are underlined and labeled. Red rectangular boxes indicate the divergent N- and C-terminal regions of MT_1_ and MT_2_.
